# Division-independent differentiation mandates proliferative competition among stem cells

**DOI:** 10.1101/208983

**Authors:** Amy Reilein, David Melamed, Simon Tavaré, Daniel Kalderon

## Abstract

Cancer-initiating gatekeeper mutations that arise in stem cells would be especially potent if they stabilize and expand an affected stem lineage (1, 2). It is therefore important to understand how different stem cell organization strategies promote or prevent variant stem cell amplification in response to different types of mutation, including those that activate stem cell proliferation. Stem cell numbers can be maintained constant while producing differentiated products through individually asymmetric division outcomes or by population asymmetry strategies, in which individual stem cell lineages necessarily compete for niche space. We considered alternative mechanisms underlying population asymmetry and used quantitative modeling to predict starkly different consequences of altering proliferation rate: a variant, faster-proliferating mutant stem cell should compete better only when stem cell division and differentiation are independent processes. For most types of stem cell it has not been possible to ascertain experimentally whether division and differentiation are coupled. However, Drosophila Follicle Stem Cells (FSCs) provided a favorable model system to investigate population asymmetry mechanisms and also for measuring the impact of altered proliferation on competition. We found from detailed cell lineage studies that FSC division and FSC differentiation are not coupled. We also found that FSC representation, reflecting maintenance and amplification, was highly responsive to genetic changes that altered only the rate of FSC proliferation. The FSC paradigm therefore provides definitive experimental evidence for the general principle that relative proliferation rate will always be a major determinant of competition among stem cells specifically when stem cell division and differentiation are independent.

**SIGNIFICANCE:** Adult stem cells support tissue maintenance throughout life but they also can be cells of origin for cancer, allowing clonal expansion and long-term maintenance of the first oncogenic mutations. We considered how a mutation that increases the proliferation rate of a stem cell would affect the probability of its competitive survival and amplification for different potential organizations of stem cells. Quantitative modeling showed that the key characteristic predicting the impact of relative proliferation rate on competition is whether differentiation of a stem cell is coupled to its division. We then used Drosophila Follicle Stem Cells to provide definitive experimental evidence for the general prediction that relative proliferation rates dictate stem cell competition specifically for stem cells that exhibit division-independent differentiation.

## INTRODUCTION

Large-scale sequencing of tumor samples, including single cells, provides information about the number and identity of mutations that drive cancer ontogeny, key initiating gatekeeper mutations and clonal histories (1–3). Understanding how each driver mutation promotes clonal selection throughout this long developmental sequence of changing cellular phenotypes and environments is very challenging, but is most approachable for the earliest mutations because they occur in the context of normal morphology and physiology. The longevity and proliferative potential of stem cells make it inevitable that the first driver mutations sometimes arise in stem cells, especially for tissues with very active stem cells and short-lived derivatives (1, 4–6). Those first driver mutations (gatekeepers) may act throughout cancer evolution but they will be especially potent if they provide a selective advantage at the earliest possible stage to stabilize a mutant lineage and amplify it to provide multiple substrate cells for sampling a variety of potential secondary mutations (6, 7). It is therefore very important to understand what types of mutations favor maintenance and amplification of an affected stem cell and hence why some gatekeeper mutations may be more potent in one tissue than another.

It might, at first thought, be expected that an increased rate of cell division would inevitably favor the amplification of any cell type. However, stem cells are generally maintained at roughly constant numbers. This constraint, generally imposed by limited space within a supportive niche environment, renders the impact of increased proliferation dependent on the strategies used for stem cell maintenance (see Figure 1) (8–10). For example, if a stem cell always divides to produce one stem cell and one differentiated cell (single cell asymmetry; model A in Figure 1), an increased rate of division of one stem cell will not alter the longevity or representation of that stem cell. Germline Stem Cells (GSCs) in the Drosophila ovary mostly undergo repeated divisions with asymmetric outcomes and mutations that alter the rate of GSC divisions do not generally affect GSC maintenance (11–14).

**Figure 1.**
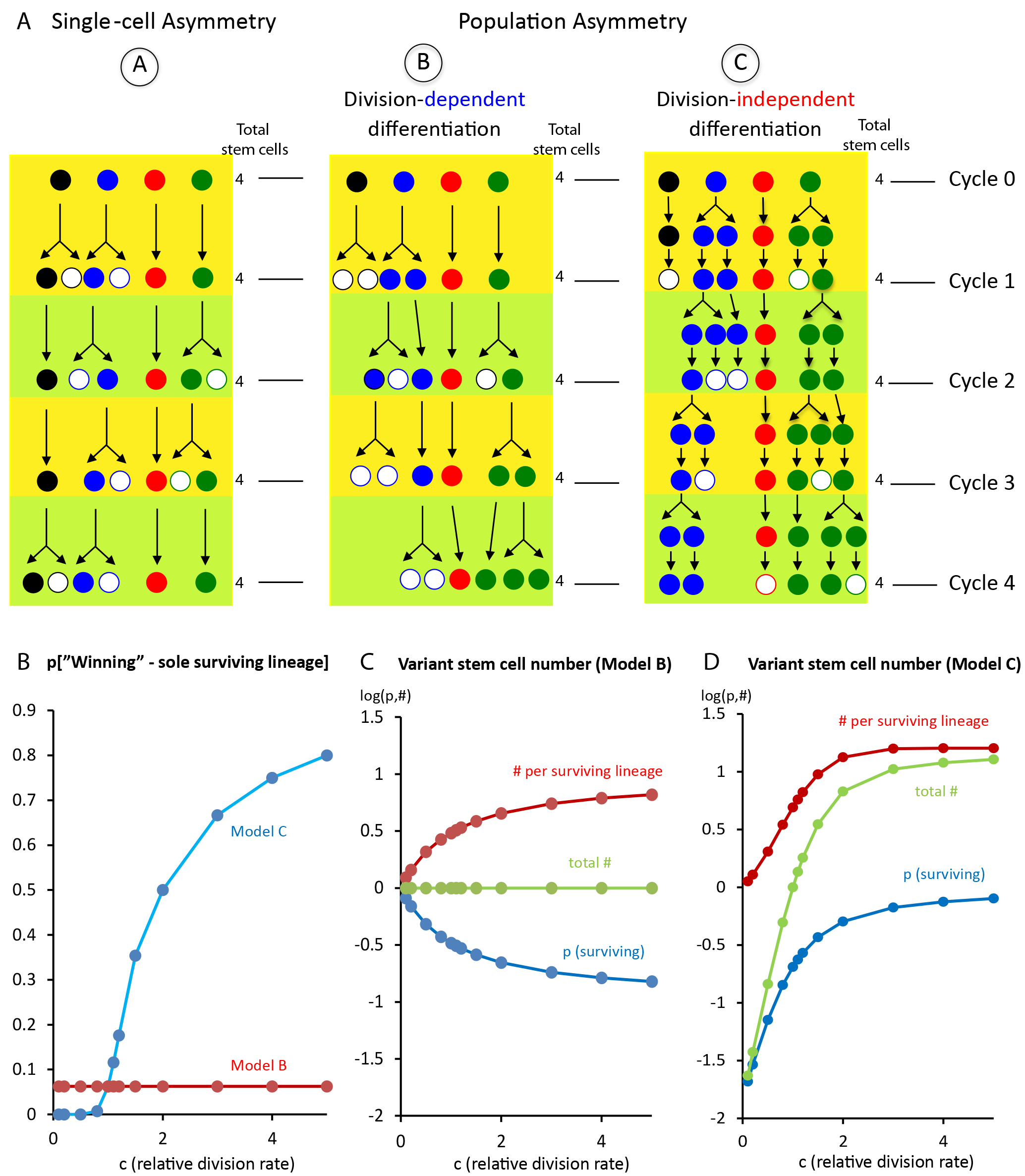
Stem cell organization dictates impact of proliferation rate on stem cell competition. (A) The cartoon depicts possible trajectories of a population of four stem cells (black, blue, red, green) through four cycles for three different types of organization. By the end of each cycle, two of the four stem cells have divided, the total number of stem cells (filled circles) remains constant (at four) and two non-stem cells (open circles) have been produced. Model A: Each stem cell division always produces one stem cell and one non-stem cell (“single-cell asymmetry”). Each lineage is therefore maintained in equal proportion no matter what their relative rates of division (here, the blue stem cells divided four times and the red stem cells did not divide at all). Models B and C represent different mechanisms of population asymmetry. Model B: Non-stem cells are only produced when a stem cell divides (“division-dependent differentiation”) but each division can produce two stem cells or two non-stem cells (with equal frequency) or one of each. The relative proliferation rate of a variant stem cell does not affect its predicted competitive success (from quantitative modeling). In this example, the red stem cell survives despite failing to divide, while the blue stem cells are extinguished despite dividing more times per cycle (4 out of 6) than even the green stem cells (3 out of 5). Model C: Non-stem cells are produced at any time, independent of division history (“division- independent differentiation”) and the total number of non-stem cells produced equals the total number of stem cell divisions over the whole population to maintain constant stem cell numbers (“population asymmetry”). The cartoon shows an intermediate stage to illustrate that division and differentiation are separate processes. Division is shown first but these processes would not be rigidly ordered. For model C, the relative proliferation rate of a variant stem cell has a large impact on its competitive success. In this example, the red stem cell is (by chance) relatively resistant to differentiation, remaining a stem cell for 3 of 4 cycles but that lineage is nevertheless extinguished eventually because the red stem cell did not divide (contrast with A and B). Conversely, even though the blue stem cells became non-stem cells at almost half of the possible opportunities (3 of 8) this lineage amplified because of frequent divisions (contrast with A and B where divisions were at least as frequent). (B-D) Graphical representation of results from quantitative modeling (Supplementary Note A) of stem cell models B and C, in each case considering a population of 16 stem cells that initially includes one variant with division frequency altered by a factor of “c”. (B) Probability that the variant stem cell lineage is ultimately the sole surviving (“winning”) lineage (constant at 1/16 for model B in red). (C, D) The probability of survival of the variant lineage (p; blue), expected number of stem cells in a surviving variant lineage (#; red) and hence (the product of p and #) the expected total number of variant stem cells present (green) for (C) model B and (D) model C, on a log_10_ scale, after a fixed time interval (this would correspond to roughly 12 cycles of egg chamber budding, or 6d, for FSCs where roughly 6 FSCs divide per budding cycle; t=12x6/16=4.5). See also Figure S1.

Several types of stem cell, including Drosophila Follicle Stem Cells (FSCs), which reside in the same ovaries as GSCs, and mammalian gut stem cells are instead maintained by population asymmetry (Figure 1). The term “population asymmetry” is generally understood to mean that the fates of two daughters of a stem cell are independent. Population asymmetry inevitably creates competition among stem cells for survival and amplification, leading to stochastic expansion of some stem cell lineages, while others are lost (“neutral competition”) (Figure S1) (15, 16). The factors that regulate competition can be uncovered experimentally by identifying hypo- or hyper-competitive genetic variants and the molecular mechanisms they affect. Both an unbiased genetic screen and analysis of a key niche signal pointed to stem cell division rate as a key determinant of FSC competition (12, 13, 17). By contrast, niche adhesion, resistance to differentiation and quiescence are more commonly cited as key parameters favoring longevity of various other stem cells, including Drosophila GSCs (9, 18). We wished to understand whether a fundamental principle of stem cell organization might explain a causal connection between proliferation and competition by using FSCs as a model stem cell.

GSCs and FSCs are housed in the germarium, which lies at the anterior of each egg-producing ovariole (Figure 2A). In the anterior half of the germarium, Escort Cells (ECs) support the differentiation of GSC derivatives into sixteen-cell cysts (19). Follicle cell precursors (FCs) then associate with germline cysts midway through the germarium and proliferate to form an expanding monolayer epithelium (20). A subset of FCs differentiate early to form polar cells and stalk cells, which allow budding of fully enveloped cysts from the posterior of the germarium to produce new egg chambers roughly every 12h, throughout the life of well-fed adult females. Until recently it was thought that each germarium contained only two, or perhaps three, FSCs, that FSCs produced only FCs and that the majority of FSC divisions produced one FSC and one FC (20-22). However, we recently reported that each germarium contains many more FSCs (about 14-16), that FSCs produce quiescent ECs as well as transit-amplifying FCs and that FSCs are maintained by population asymmetry (23).

**Figure 2.**
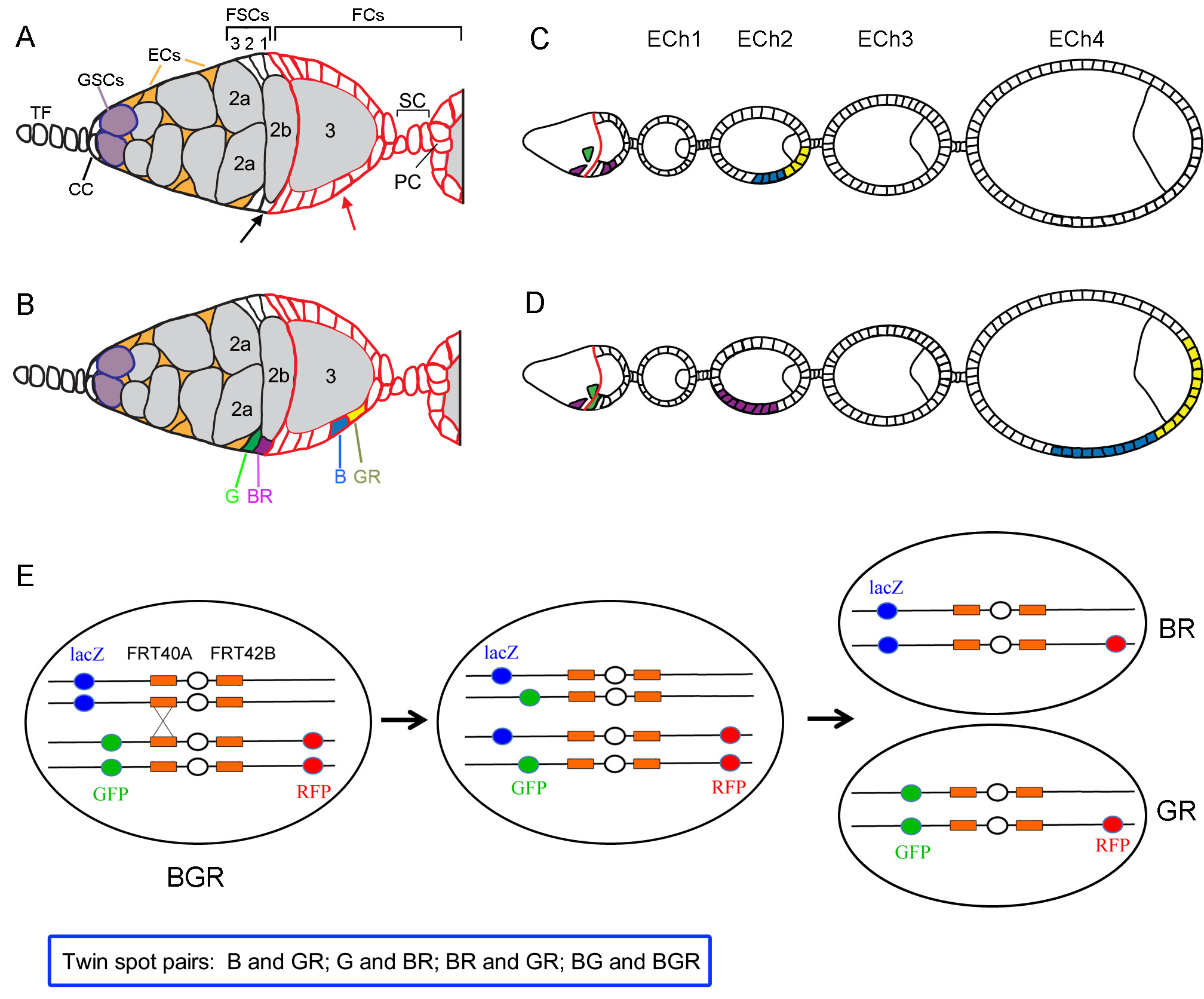
Drosophila oogenesis and twin-spot analysis of FSC daughter fates. (A-D) Illustration of FSC and FC twin-spot clones. (A) Germarium diagram showing Terminal Filament (TF) cells, Cap Cells (CC), Germline Stem Cells (GSCs), GSC daughters developing into 16-cell germline cysts (light grey), Escort Cells (EC, orange), Follicle Stem Cells (FSC) and Follicle Cells (FC), including Stalk Cells (SC) and Polar Cells (PC) from anterior (left) to the newest egg chamber. Fas3 expression on FC surfaces is shown in red. The anterior limit of Fas3 staining, running along the posterior surface of a stage 2b germline cyst, provides a key landmark. FSCs lie in three layers (“3-1”) immediately anterior to Fas3 but posterior to 2a cysts. (A-D) illustrates the progression over time of the products of mitotic recombination in an FSC (black arrow) and an FC (red arrow). (B) Germarium showing twin-spot daughters immediately after recombination in an FSC (green, G and purple, BR) and in an FC (blue, B and yellow, GR). Letters indicate the presence of a given transgene (B- Blue *lacZ*, G-Green GFP, R- Red, RFP). (C, D) The B and GR FC daughters proliferate to form patches, which are always on the same egg chamber, as it grows and moves to the posterior (right) along the ovariole (C) two cycles (24h) and (D) four cycles (48h) after initial marking. Egg chambers bud from the germarium roughly every 12h. (C) A BR FC produced in the previous cycle has divided once, leading (D) to an FC patch on the second egg chamber two cycles later. Unpaired FC patches, as shown here for BR, must derive from recombination in an FSC and were never observed beyond the fourth egg chamber 72h after heat-shock. (E) The starting genotype at the time of mitotic recombination is shown (left) for the second chromosome of flies used for twin-spot lineage marking. The *tub-lacZ* (“lacZ”), *ubi-GFP* (“GFP”), *ubi-RFP* (“RFP”) transgenes, as well as *FRT 40A* and *FRT 42B* recombination targets (orange) either side of the centromere (white oval) are indicated. Heat-shock induction of a *hs-flp* transgene on the X-chromosome can induce (middle panel) recombination at either or both pairs of homologous FRTs, followed by (right panel) segregation to yield two daughter cells with recombinant genotypes in predictable twin-spot pairings (here BR and GR daughters are produced; other possible pairings are B:GR, G:BR and BG:BGR).

We first considered two potential mechanisms for population asymmetry (models B and C in Figure 1) from a theoretical perspective and used quantitative modeling to conclude that the impact of proliferation rate on stem cell competition should depend critically on whether differentiation is independent of stem cell division. We then examined FSC organization in more detail, specifically to discover whether FC production was temporally coupled to FSC division and to test rigorously the impact of mutations that altered FSC proliferation rate on FSC representation. The experimental results provide definitive evidence for the general principle that stem cell competition depends on relative proliferation rates specifically when stem cell division and differentiation are independent processes.

## RESULTS

### Contrasting impacts of altered proliferation for different population asymmetry mechanisms

We considered three idealized strategies for stem cell maintenance to evaluate from a theoretical standpoint how stem cell organization controls the impact of cell proliferation rates on stem cell competition. If each stem cell division produces an asymmetric outcome (model A, Figure 1A), there will be no competitive advantage or disadvantage for a stem cell that divides at a different rate. Drosophila ovarian GSCs appear to show this organization and indifference to stem cell division rates (12, 13).

For stem cells governed by population asymmetry there are two possible underlying mechanisms that have not generally been explicitly distinguished experimentally or conceptually. The predicted consequences of altered proliferation are widely different for the two models. If stem cell division and differentiation are rigidly coupled (model B, Figure 1A), then an individual stem cell that proliferates faster than others (blue and green cells in Figure 1A) will have a higher chance of amplification during a fixed time interval, but it will also have a proportionally higher chance of being lost. Hence, a qualitative appraisal suggests there will be little or no net consequence on stem cell competition. This organization is sometimes assumed and might occur, for example, if stem cells are maintained by niche adhesion and only release a rigid attachment to niche cells at the time of cell division (24, 25).

If stem cell division and differentiation are independent processes that are not linked mechanistically or temporally (model C, Figure 1A), then each stem cell division initially produces two stem cells and a stem cell can differentiate at any time. Here, an increase in proliferation rate of one stem cell relative to others will inevitably lead to a higher likelihood of amplification and a reduced likelihood of losing the variant lineage (Figure S1). Conversely, stem cells that rarely divide (red cells in Figure 1A) can survive for long periods if differentiation is coupled to cell division (models A and B) but are very likely to be lost within a few cycles if differentiation is independent of stem cell division (model C) because there is a chance to differentiate at every cycle (time period). In summary, there is a strong likelihood that slower proliferating stem cells will be lost and faster proliferating stem cells will amplify only when there is division-independent differentiation (model C).

The different organizations described above for population asymmetry were translated into a quantitative model in order to evaluate whether an altered proliferation rate has any effect on stem cell competition in model B and to predict the magnitude of such effects in model C (see Supplementary Note A and Figure S1). In each case, the model was constrained to maintain a constant total number of stem cells. Thus, if an extra stem cell is produced at any time (by a division producing two stem cells), this was immediately followed by stem cell loss (by a division producing two non-stem cell daughters in model B or by differentiation in model C) and, conversely, stem cell loss was followed by stem cell duplication. Additionally, the probabilities of a division yielding two stem cells or two non-stem cells were considered to be equal in model B. These models can be treated as classical Markov chains (see Supplementary Note A for details).

For model B, a variant stem cell with an altered division rate (by a factor, c) has an unchanged probability of being the sole lineage remaining after all others have been lost (Figure 1B, Figure S1 and Supplementary Note A). By contrast, in model C the probability of being the winning (sole remaining) lineage increases greatly for a variant stem cell that divides faster than its competitors (Figure 1B and Supplementary Note A). For a population of 16 stem cells, a 50% increase in proliferation rate raises the probability of indefinite survival from 1/16 to about 1/3 (more than 5-fold) and reduces the expected average time to achieve that state two-fold (Supplementary Note A).

The average number of stem cells in a lineage initiated from a single variant stem cell at any time prior to completion of clonal evolution (Figure S1) is predicted to be the same for all values of c in model B (Figure 1C, green line). That is because changes in the average number of stem cells per surviving variant lineage are exactly offset by inverse changes in the probability that the variant lineage survives (Figure 1C). By contrast, both of these parameters increase for greater c values in model C and therefore sum to give an even larger response for the total number of variant stem cells present (1.0 (c=1.0), 1.8 (c=1.2), 3.5 (c=1.5) and 6.7 (c=2.0) expected variant stem cells; Figure 1D). In summary, quantitative modeling shows that for model B an altered rate of proliferation has no impact on cell competition measured by the eventual winner of clonal competition or stem cell representation at earlier times, whereas altered division rates have a large effect on both measures of competition for stem cells organized as in model C.

### Twin-spot lineage analysis to reconstruct FSC behaviors over 72h

To test the theoretical predictions connecting stem cell organization to the impact of altered proliferation on competition, which should apply to all types of stem cell, we turned to Drosophila FSCs. To determine whether FSC differentiation into FCs occurs only at the time of FSC division (model B) or independent of FSC division (model C) we tried to reconstruct the precise behavior of FSCs over a 72h period through a detailed lineage analysis. Marked clones were created at a fixed time by using a heat-shock induced *flp* recombinase to promote mitotic recombination at *FRT* sites located at the base of chromosome arms harboring *GFP* (“G”), *β*-galactosidase (“B”) and *RFP* (“R”) transgenes (Figure 2E). We showed previously that in these flies fewer than one in a hundred ovarioles produced recombinant FSC genotypes in the absence of heat-shock (23). Hence, we can be sure that virtually all recombination events occur shortly after heat-shock.

Our first objective was to define the first egg chambers populated by FC-derivatives of recombination in an FSC. Both FSCs and their proliferative FC progeny can undergo mitotic recombination to produce twin-spot daughters with predictable pairs of color combinations (B:GR, G:BR, BR:GR, and BG:BGR daughter pairs) (Figure 2E). However, the earliest FCs are distinguished from FSCs by their association with a developing germline cyst, leading to the inevitable passage of an FC and all of its progeny through the ovariole. Recombination in an FC therefore always produces two daughters associated with the same germline cyst; those daughters will then proliferate to form paired twin-spot FC patches on the same egg chamber (illustrated for B:GR FC daughters in Figures 2A-2D). By contrast, an FC patch that has no paired twin-spot on the same egg chamber must have derived from recombination in an FSC (illustrated for G:BR FSC daughters in Figures 2A-2D).

At 72h after heat-shock, unpaired FSC-derived FC patches were found in the fourth youngest egg chamber for more than a quarter of all ovarioles; older egg chambers always contained only paired twin-spot FC patches. We therefore deduced that the first opportunity for a marked FSC daughter to become an FC in any of the experimental ovarioles was the passage through the FSC region of a germline cyst that will become the fourth youngest egg chamber 72h later (as illustrated for the BR FSC daughter in Figure 3D). This deduction is consistent with the expectation that egg chambers bud from the germarium roughly every 12h (20) and that the germarium generally contains two cysts posterior to the FSCs, leading to a maximum of six cycles of FC recruitment over 72h (Figure 3D). Egg chamber production in all ovarioles of the experimental flies was likely synchronized to within 12h. We therefore made the conservative assumption that marked FSC daughters had the opportunity to contribute to all egg chambers up to the third youngest (five cycles of egg chamber budding in total) for ovarioles with no unpaired FC patches in the fourth youngest egg chamber (as in Figure S3).

**Figure 3.**
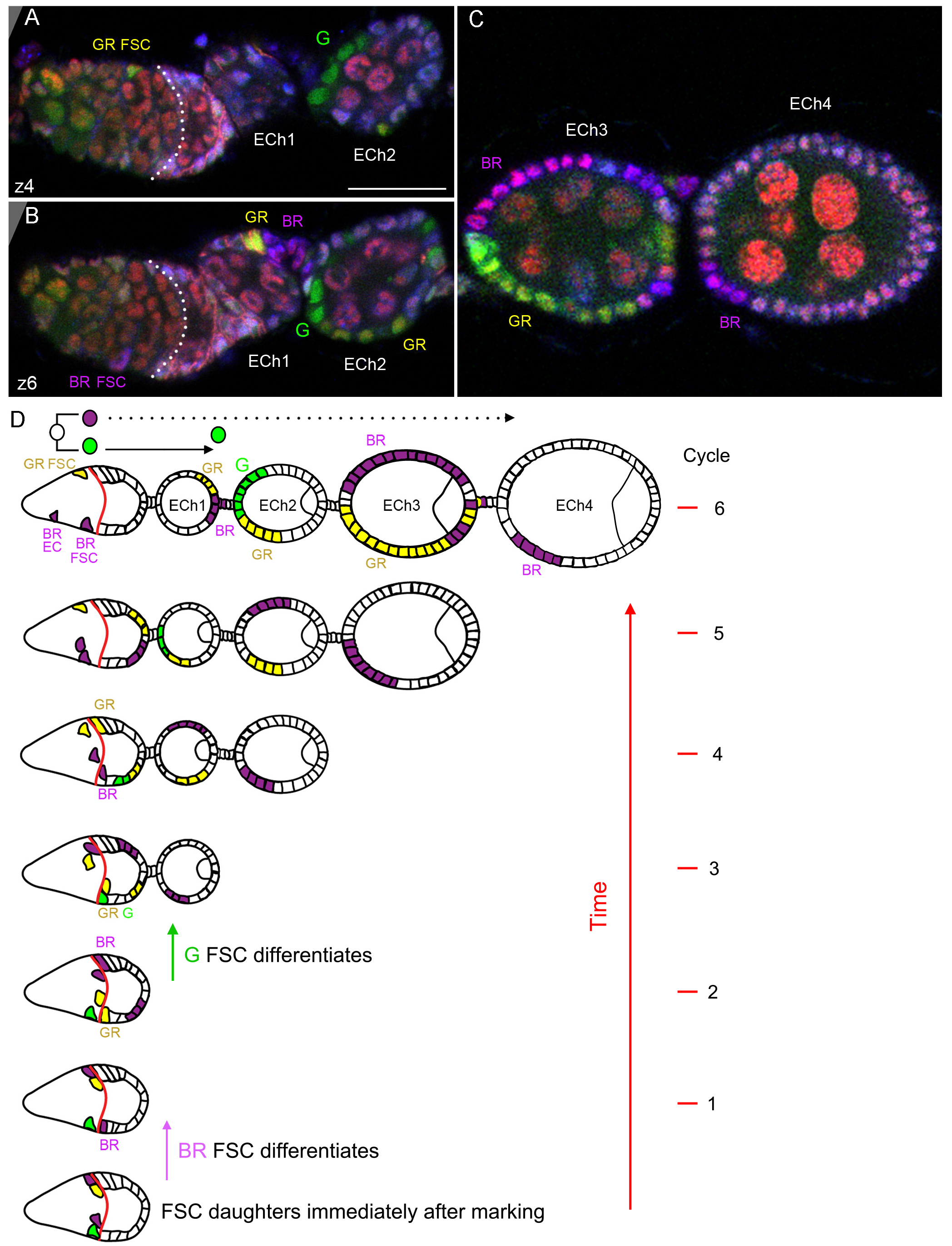
Twin-spot lineage analysis to determine when an FSC daughter becomes an FC. (A-D) Analysis of FSC twin-spot clones 72h after induction. (A, B) Two z-sections of the germarium and first two egg chambers (“ECh”) and (C) egg chambers three and four of an ovariole illustrated in the top cartoon of (D). All z-sections were examined in each fluorescence channel to assign all cell colors definitively (see Figure S2). The anterior (left) limit of surface Fas3 staining (pink) is outlined with white dotted lines. Scale- bar is 20lm. (A-D) The BR (purple) FC patch in egg chamber 4 had no matching twin-spot FCs (would be G or GR) in the same egg chamber and must therefore derive from recombination marking in an FSC. Hence, all marked cells up to and including ECh4 derive from daughters of recombination in an FSC. The colors present up to ECh4 are G, BR and GR, showing that one FSC produced a G:BR twin-spot pair (illustrated above the ovariole in (D)) and a second FSC produced a GR:BR twin-spot daughter pair. There is only one G FC patch, no G FSC and no G ECs, implying that there were no divisions of the G daughter of the FSC after it was born (shortly after the time of heat-shock). The G FC patch is in ECh 2 and was therefore produced two cycles (about 24h) after the first opportunity to become an FC (in ECh 4, as for the BR FC). Thus, the G daughter of a FSC became a FC long after it was born (about 24h; two cycles of egg chamber budding). (D) The inferred histories of the G:BR and BR:GR twin-spot pairs are illustrated from immediately after recombination marking (bottom) through each 12h cycle of egg chamber budding up to the final stained ovariole at 72h after heat-shock. All FCs produced from cycle 1-5 are labeled at the cycle produced and in the final ovariole cartoon.

### Division-independent differentiation of FSCs to become FCs

For each lineage derived from mitotic recombination in an FSC we could see how many FSCs remained after 72h, whether any ECs had been produced and exactly when any FCs had been produced because the order of egg chambers displays the time at which a founder FC associated with a passing germline cyst (Figure 3D). We looked for examples of an FSC daughter lineage that included only a single patch of FCs, no ECs and no FSCs. That pattern reports a daughter of mitotic recombination in an FSC that became a founder FC without any intervening divisions; it is exemplified by the G lineage in Figure 3 (single channels shown in Figure S2) and the B lineage in Figure S3 (also see Supplementary Note B and Figure S4). We found seventeen such examples. In six cases, the solitary FC patch was in the fourth youngest egg chamber (egg chamber 4), implying that the marked cell became an FC immediately, or shortly after the FSC division where mitotic recombination occurred. In three cases, the FC patch was in the third youngest egg chamber. In these three examples, we cannot be certain if FC production was immediate or delayed by one cycle because the germline cyst that became the fourth youngest egg chamber may also have been available for population after the marked FSC daughters were born. Importantly, in eight cases (including the G lineage in Figure 3, and the B lineage in Figure S3) the FC patch was in egg chamber 2 or younger; moreover, an older egg chamber contained FCs from a different marked FSC derivative. In these eight cases, we can deduce that a marked FSC daughter was born shortly after heat-shock, did not divide subsequently, and then became an FC only after one or more cysts had passed through the FSC region. In other words, those FSC daughters only became FCs 12-60h after birth (Figures 3D and S3D). We repeated the experiment, examining a new set of ovarioles 72h after heat-shock and found a similar distribution of locations for solitary marked FC patches (22/35 prior to egg chamber 3). These observations provide direct evidence that an FSC can become an FC at any time, not just immediately after cell division. Thus, FSCs exhibit “division-independent differentiation” and conform to model C (Figure 1A).

We also examined all ovarioles harboring only one pair of twin-spots that likely derived from a single stem cell (see Supplementary Note C) to determine the immediate behavior of FSC daughters. If a marked daughter produced two or more cells (ECs, FC patches or FSCs) by 72h we deduced that it must have divided as an FSC before any subsequent differentiation event (GR FSC in Figure 3; BR FSC in Figure S3). In nine cases, both FSC daughters divided as a stem cell prior to any further events; in two instances, both daughters became FCs without any intervening divisions; in two cases, one daughter divided as a stem cell while the other became an FC (one example) or an EC (one example) without an intervening division. Altogether, 20 FSC daughters subsequently divided as a stem cell, while five became FCs and one became an EC without dividing again as an FSC. These outcomes are consistent with a key aspect of division-independent differentiation (model C, Figure 1A), namely the initial production of two stem cells from all FSC divisions.

### Genetic evidence for division-independent differentiation

As a further test of whether or not production of FCs is contingent on concurrent FSC division, we examined genetic conditions that reduced the rate of FSC division substantially. We previously identified loss of function mutations affecting the cell cycle regulator, Cyclin E (CycE) and a DNA replication component, Cutlet, as reducing FSC maintenance (12, 13). We also showed that loss of Yorkie (Yki) activity reduced FSC proliferation and FSC maintenance (17). Here, we found that the rate of FSC division, measured by EdU incorporation 6d after clone induction, was indeed greatly reduced relative to controls for *cycE* (17%), *cutlet* (34%) and *yki* (11%) mutant FSCs (Figure 4E) marked as GFP-positive by the MARCM technique (26). To measure FC production we examined all ovarioles with a marked FSC and counted the proportion of germarial cysts and egg chambers that included a marked FC patch (Figure 4). All three proliferation-defective mutant FSC genotypes produced substantial numbers of FC patches (Figure 4). Indeed, the proportion of cysts and egg chambers with marked FCs per marked *cycE*, *cutlet* or *yki* FSC was 11.5% (combining all three genotypes), only marginally lower than for controls (average: 14.5%). Thus, drastic reduction of FSC division frequency did not significantly curtail FC production, providing further evidence that FCs can be produced at any time, not only when an FSC divides.

**Figure 4.**
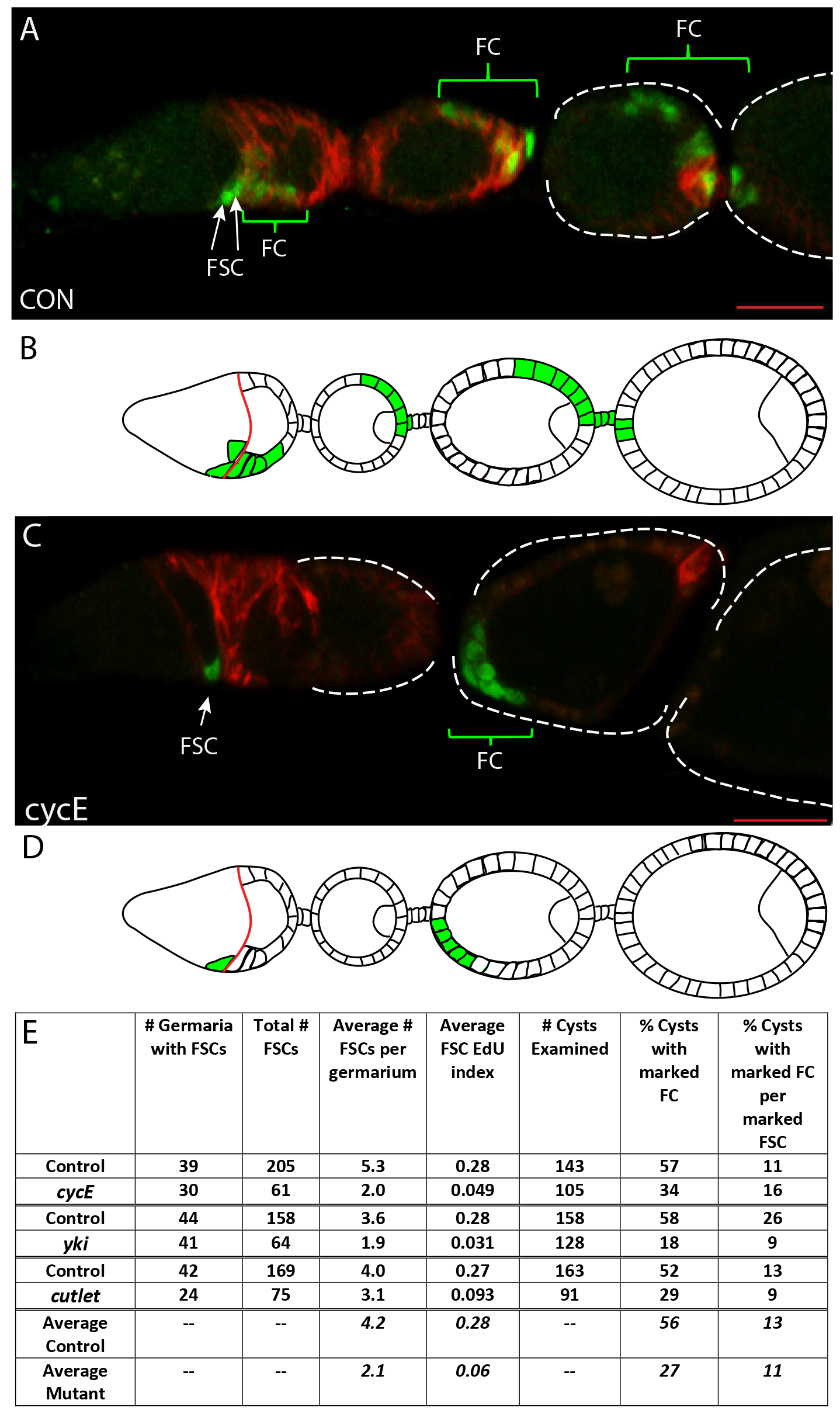
Proliferation-deficient FSCs still produce FCs. (A-D) Ovarioles with MARCM clones for (A) control and (C) cycE^WX^ genotypes, labeled with Fas3 (red) 6d after clone induction. Green MARCM-labeled FSCs are within three cell diameters to the left (anterior) of the Fas3 staining border (white arrows). FC patches in the germarium or egg chambers (outlined with dashed white lines where Fas3 staining outline is weak) are indicated by brackets. Scale-bar is 20am in each case. (B, D) Schematics of ovariole images above, showing the locations of marked FSCs and FCs. (E) Summary of FSC division rates (EdU index), FSC numbers, FC patches and FC patches per FSC for mutants with reduced FSC proliferation together with their controls.

### Reconstructing FSC histories to detail FSC dynamics

We also used the detailed record of FSC behavior manifest by twin-spot clones to confirm and extend previous conclusions about FSC numbers, FSC dynamics and FC production. In the two twin-spot experiments, the average percentage contribution of a single color to the FCs of an egg chamber for B, G and BG clones was 18.9% (18.9% and 18.8% in the two sets of experiments) and for solitary FC patch clones (where single FC founders are almost certain) it was 17.8% (19.0% and 16.5%). Hence, our best estimate of the average number of founder FCs per egg chamber is between five and six (1/0.189 = 5.3, 1/0.178 = 5.6). We also measured the early rates of FSC loss and FSC amplification. We found that 25 of the 49 marked FSC daughters likely arising from a single cell were lost (becoming FCs or ECs) over the next 3d. This high rate of loss supports a model of population asymmetry, where individual stem cells are frequently lost or amplified in a stochastic process of neutral competition.

Finally, we derived explicit histories of FSC behavior for all marked FSC daughter lineages in order to calculate the average frequency of FSC divisions and the average frequency of differentiation to FCs and ECs. To facilitate modeling, and in keeping with our deduction of division-independent differentiation, we artificially split each cycle of egg chamber budding into an opportunity for all FSCs to divide, followed by an opportunity for all FSCs to become an FC or EC (as in model C of Figure 1A). The stained ovarioles at 72h showed the total number of FSCs, FC patches and ECs produced by each lineage as well as the cycle at which founder FCs were produced (Figure 3 and Figure S3). The cycles at which marked FSCs divided were either definitively compelled or highly constrained by the sequence of FC production together with the total number of FSCs and ECs produced. Wherever FSC divisions could equally likely have occurred at either of two different cycles, assignments were made so that FSC divisions were spaced as evenly as possible (see Figure S3D legend).

By combining the cycle-by-cycle inferred histories of 79 lineages (illustrated and tabulated for one ovariole in Figure S3), we found that marked FSCs divide at 44% (221/501) of available opportunities (each cycle represents an opportunity for each FSC) and that marked FSCs become FCs at 21% (159/747) of available opportunities, while producing 43 ECs over the same period (1 EC per 3.7 FCs). If each egg chamber is seeded by five founder FCs, then 1.4 (5/3.7) ECs are produced at each cycle on average, and a total of 6.4 FSC divisions would therefore maintain homeostasis. For 6.4 divisions at an average frequency of 0.44 per FSC there must be, on average, 14.5 (6.4/0.44) FSCs at the start of a cycle. Similarly, to produce 5 FCs per cycle at the frequency observed (0.21 per FSC) there should be 23.8 FSCs (5/0.21) in the middle of a cycle. At that stage the number of FSCs is artificially inflated by 6.4 in our model because FSCs have divided but none has become an FC or EC. So, the true estimate of the average number of FSCs based on FC production rate is 17.4 (23.8-6.4). These two estimates (14.5 and 17.4, based on FSC division and FC production frequencies, respectively) are in good agreement with the earlier estimate of 14-16 FSCs based on counting the number of surviving FSC lineages over time and counting the total number of cells within the FSC domain (23). Thus, our analysis of twin-spot FSC lineages has confirmed our recent conclusions about FSC numbers and FSC maintenance by population asymmetry, it has revised our best estimate of the number of FC founders per egg chamber and demonstrated that differentiation of an FSC to an FC is not dependent on FSC division.

### FSC competition is dictated by relative rates of proliferation: experimental evidence

Division-independent differentiation of FSCs predicts that competition amongst FSCs will be determined substantially by their relative rates of proliferation (Figure 1). There is already substantial evidence that FSC proliferation rate strongly influences FSC competition (12, 13, 17). However, previous analyses of competition between FSCs was limited to measuring the loss of a marked variant FSC lineage over time and, for the rare changes that enhanced competitive success, by counting the proportion of ovarioles containing “all-marked” clones, where a single lineage contributes all FCs to several successive egg chambers (12, 17). The recent findings, confirmed here, that each germarium contains many FSCs (14–16) and that the number of FSCs in each marked lineage changes over time as a result of competition (23), allow a better measure of stem cell competition as the average number of FSCs present at a fixed time after FSC clone induction (Figure S5).

Here we measured FSC proliferation rates according to EdU incorporation over one hour of in vitro incubation immediately after ovary dissection (Figure 5B-I). We measured FSC competition (how well a variant stem cell survives and amplifies within a niche containing a constant total number of stem cells) by counting the average number of marked FSCs per ovariole 6 and 12d after clone induction for a variety of FSC clone genotypes expected to affect proliferation (Figure 5B-M and Figure S5). FSC clones with a homozygous mutation or expressing a GAL4/UAS- driven transgene were generated and marked as GFP-positive by the MARCM technique (26). These clones were compared to control FSC clones generated in the same way and strictly in parallel in flies lacking the mutation or transgene under investigation.

**Figure 5.**
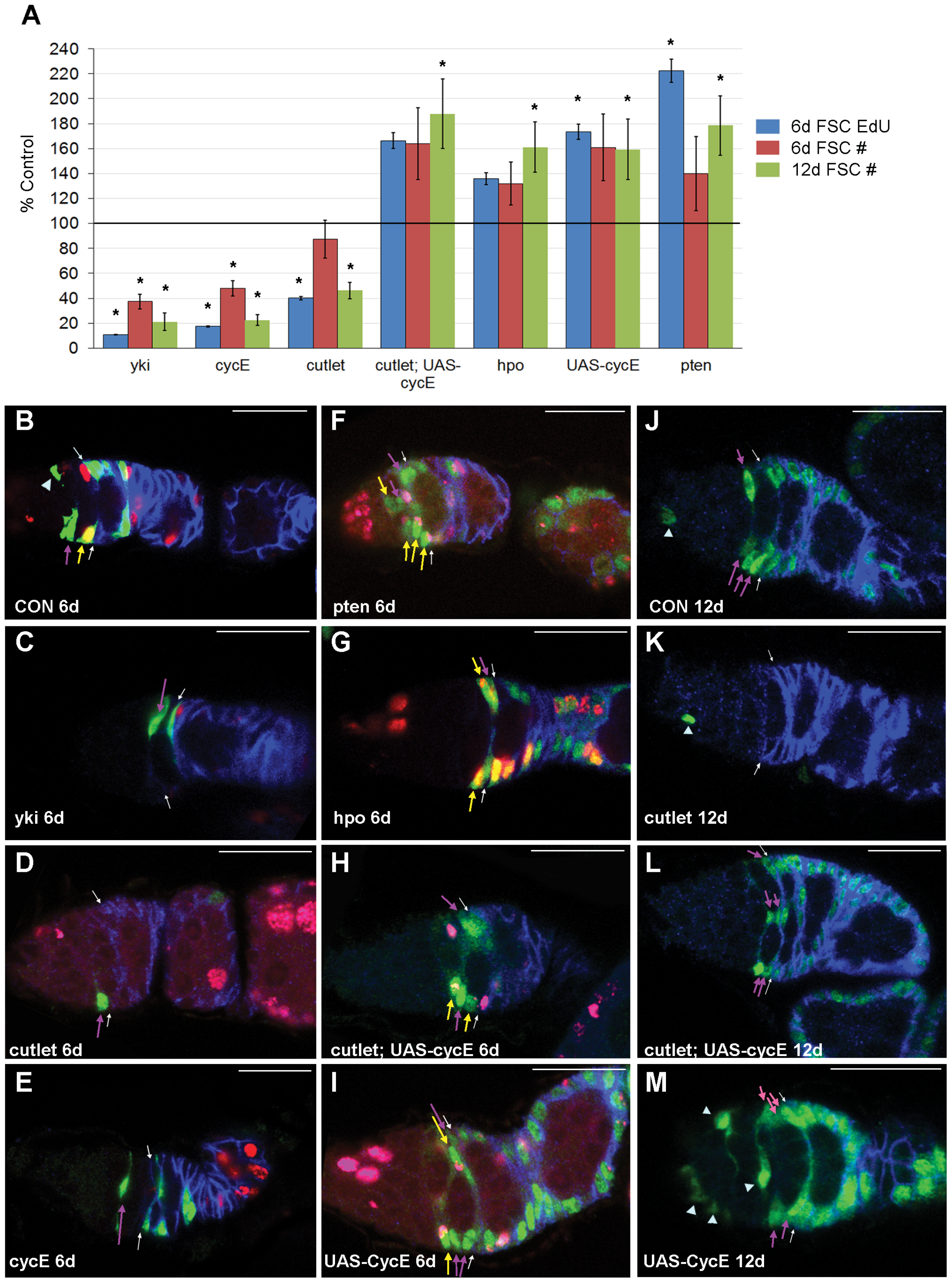
FSC competition is determined by relative FSC proliferation rates. (A) Correlation between proliferation rate (blue: FSC EdU index) at 6d and average number of marked FSCs per ovariole at 6d (red) and at 12d (green), expressed as percentage of control values for MARCM FSC clones of the listed genotypes. Error bars show SEM (EdU: n= 64 (158), 61 (205), 75 (159), 173 (159), 201 (158), 193 (159), 141 (159) FSCs in the order shown, 6d FSC#: n= 58 (54), 38 (61), 35 (65), 43 (65), 52 (54), 49 (65), 41 (65) ovarioles in the order shown, 12d FSC#: n= 58 (69), 53 (55), 37 (71), 56 (71), 63 (69), 56 (71), 54 (71) ovarioles in the order shown; values in parentheses are for the associated controls). Significant differences to control EdU index (by Fisher’s exact two-tailed test, p<0.05) and control marked FSC number (by Student’s t-test, p<0.05) are indicated. (B-M) Representative germaria from MARCM analyses summarized in (A). (B-I) Germaria with MARCM clones of the designated genotypes (green), labeled with EdU (red/pink) and Fas3 (blue) 6d after clone induction. (J-M) Germaria with MARCM clones (green), labeled with Fas3 (blue) 12d after clone induction. (B-M) Green MARCM-labeled FSCs with EdU (yellow arrows; green and pink often adjacent in same nucleus rather than overlaid) or without EdU (purple arrows) are within three cell diameters to the left (anterior) of the Fas3 staining border (white arrows); ECs (arrowheads) are further anterior. At 6d, (B-E) there are fewer labeled FSCs for *yki*, *cutlet* and *cycE* than for controls and lower frequencies of EdU (red) incorporation into the labeled FSCs. (F-I) By contrast, the number of labeled FSCs and the frequency of EdU incorporation is higher for *hpo*, *pten* and both genotypes that include *UAS-cycE*. (J-M) By 12d, the number of marked FSCs has declined further for (K) *yki*, *cycE* and *cutlet* and remains higher for (L, M) *cutlet; UAS-cycE* and *UAS-cycE* relative to controls. Scale-bar is 20rm in each case. See also Figure S5.

We found that *yki*, *cycE* and *cutlet* mutations reduced FSC proliferation and also substantially reduced the average number of marked FSCs per germarium at 6d and 12d (Figures 5A-E, J, K). Moreover, expression of excess CycE restored both the proliferation rate and the average number of marked *cutlet* mutant FSCs to levels above those of controls (Figure 5A, D, H, K, L). Conversely, excess CycE alone, loss of *hpo* (which increases Yki activity), or increased PI3 kinase pathway activity due to mutation of *PTEN* (12) increased both the proliferation rate of marked FSCs and the average number of marked FSCs per germarium (Figure 5A, B, F-J, L, M).

It is possible that changes in the activity of Cutlet, the PI3 kinase pathway, Yki or even CycE may have affected FSC competition by changing a property other than proliferation rate. Alterations to Wnt signaling provide a precedent for changes that affect FSC numbers by altering the likelihood of FSC differentiation. Increased Wnt signaling caused FSC loss due to excessive conversion into ECs, whereas loss of Wnt signaling increased the likelihood of conversion into FCs, which was measured by the accumulation of FSCs in the most posterior, FC-adjacent FSC layer (“layer 1”) and by the proportion of FSC clones associated with FCs (23). We therefore measured these parameters for the genetic changes we used to alter FSC proliferation rates.

Genetic changes that reduced marked FSC numbers did not increase the number of marked ECs produced, the proportion of marked FSCs in layer 1 or the proportion of marked FSC clones with marked FCs, ruling out enhanced differentiation to ECs or FCs as responsible for the FSC deficit (Figure S6A-D). Conversely, these measures of marked EC and FC production were not decreased by alterations that increased marked FSC numbers (Figure S6A, B). Indeed, slower-dividing FSC variants were slightly biased towards anterior layers and the number of marked ECs present generally correlated positively with the number of marked FSCs, consistent with changes in EC production simply following a primary change in the number of their progenitor FSCs. These observations fully support the conclusion that the drastic decreases (*yki*, *cycE*, *cutlet*) or increases (*hpo*, *UAS-cycE*, *pten*) in marked FSC numbers we observed were caused by changes in the rate of proliferation *per se*, rather than by any unanticipated, secondary effects on FSC location or differentiation (Figure 5A and Table S1). Thus, FSCs provide robust direct evidence for a general model of organization of stem cells, namely population asymmetry with division-independent differentiation, where relative proliferation rate is both predicted and shown experimentally to be a key determinant of which stem cells are the most competitive.

## DISCUSSION

We have followed the behavior of individually marked FSCs in detail to show that FSC differentiation is not coupled to FSC division. This organization represents a subset of population asymmetry models and predicts that stem cell proliferation rate will be a major determinant of stem cell competition. We confirmed prior strong evidence of a causative link between proliferation rate and competition among FSCs (12, 13, 17) still more rigorously by measuring the proliferation rates and competitive outcomes for a number of genetic alterations that appear only to affect proliferation. The important general implication of these findings is that an analogous organization of any stem cell population, defined by the key characteristic of division-independent differentiation, will necessarily render those stem cells prone to cancer-promoting gatekeeper mutations that increase the rate of stem cell proliferation.

### Stem cell dynamics constrained by niche space

Stem cells generally require a specific environment to be maintained. If that requirement limits the space where stem cells can survive, then a specific stem cell lineage can only expand at the expense of others; it cannot expand independently or indefinitely (18, 27). This constraint applies to the normal FSC niche, to our theoretical modeling and, for example, to mammalian intestinal stem cells in a single crypt; it is a key reason why only one category of stem cell organization (Figure 1A) permits a causal connection between differential proliferation and competition.

Some mutations that alter stem cell proliferation might additionally relieve, or substitute for required niche factors and therefore allow the entire stem cell domain to expand. Those mutations could be particularly potent primary changes leading to expansion of a stem cell lineage within a single niche or they could lead to a secondary expansion of a lineage, as in the accelerated colonization of neighboring intestinal crypts (28, 29). Those consequences would not be limited to stem cell populations exhibiting division-independent differentiation but the effects on stem cell competition would also not be due solely to a change in stem cell proliferation rate. For FSCs, strong hyper-activation of JAK-STAT signaling appears to expand the FSC domain dramatically (30); the genetic changes studied in this work did not show any clear evidence of altering the FSC domain.

### FSCs and mammalian intestinal stem cells as archetypes of proliferation-dependent competition

It has generally not been possible to follow endogenous stem cell behavior in enough detail to determine whether stem cell differentiation is coupled to cell division. The singular, notable exception prior to our work is a live imaging study of mammalian intestinal stem cells. That study revealed conversion of stem cells to transit-amplifying (TA) cells without very recent cell division (31), just as we observed for FSCs. In fact, many aspects of the overall organization of FSCs and mammalian intestinal stem cells are remarkably similar. These include the size of the stem cell population, rapid constitutive stem cell divisions and now, division-independent differentiation (23, 32, 33).

Activating mutations in the Wnt or Ras pathways that increased the rate of mouse intestinal stem cell proliferation were found to promote stem cell survival and amplification, though it was not explicitly tested whether this outcome was dependent on altered proliferation rather than additional effects of those pleiotropic pathways, such as directly modulating differentiation (34, 35). In fact, Wnt signaling is known to affect intestinal cell locations and the nature of stem cell products, while Ras activation was also shown to increase the rate of crypt fission, effectively expanding the niche for an otherwise spatially constrained stem cell population (7, 34). Despite these reservations about experimental proof of a causal connection, we can confidently predict that intestinal stem cells must, indeed, exhibit a strong influence of proliferation rate on stem cell competition specifically because they undergo division-independent differentiation. This connection was not previously highlighted as causative or fundamental (31). Our study of FSCs explicitly spells out this important, universally applicable connection and provides robust experimental evidence that stem cell competition is guided by relative proliferation rates for stem cells that undergo division-independent differentiation, as described below.

Previously, the major niche signal, Hedgehog (Hh) was shown to regulate FSC competition principally by transcriptionally inducing the co-activator Yorkie (Yki), and Yki was shown to act by inducing CycE to induce proliferation (17). Here we showed that alteration of Yki activity and additional manipulations of factors expected to alter only proliferation (CycE, Cutlet), as well as changes to PI3 kinase activity, produced corresponding changes in FSC proliferation rate and FSC numbers; fewer FSCs in response to reduced proliferation and more FSCs when proliferation rates were higher. Moreover, other potential causes of the observed changes in FSC numbers (FSC location and the rate of conversion of FSCs to ECs or FCs) were ruled out by directly measuring these parameters. Hence, the cumulative experimental evidence linking stem cell proliferation rate to competition is currently stronger for FSCs than for any other stem cell (36). Moreover, the consequences of activating mutations in the Hh or Hpo/Yki pathways in FSCs provides a clear paradigm for how a gatekeeper mutation affecting a signaling pathway that controls stem cell proliferation can lead to pre-cancerous amplification of an affected stem cell (17, 37).

### Proliferation-dependent competition and stem cell exhaustion; different time-scales or stem cells?

Our studies concern a relatively short time-frame that is plausibly relevant for the amplification of a stem cell harboring a primary mutation that could eventually lead to cancer. Some mutations that increase proliferation and lead to stem cell amplification in the short term might also eventually have a deleterious effect on stem cell survival, perhaps because of DNA damage from excessively fast or incessant replication. The latter possibility, sometimes termed “stem cell exhaustion” is often cited for hematopoietic stem cells (HSCs) and provides an attractive general rationale for minimizing the normal replicative duties of at least a subset of stem cells, as observed experimentally for HSCs (38–40). Intestinal crypts also contain relatively quiescent stem cells that can replace the population of actively dividing stem cells in emergency situations (32). Amongst normal FSCs in a germarium we have also observed spatial heterogeneity of proliferation rates and it is not yet known whether quiescent ECs might become FSCs under normal or stress conditions (23).

For HSCs, many, but not all, genetic changes that increased proliferation rate led to a long-term reduction in HSC potency measured by a transplantation assay, while HSC function over the short term and under physiological conditions was not measured (38, 39, 41). The organization of HSC niches and HSC dynamics are also not sufficiently well understood at present to know whether differentiation depends on stem cell division. Consequently, the relevance of the concepts discussed in this work to normal HSCs and early steps in blood cancers is not excluded by earlier conclusions of proliferative stem cell exhaustion and remains to be explored. Conversely, while further studies are warranted, we are not aware of significant evidence for proliferative exhaustion of FSCs or mammalian intestinal stem cells (15, 16, 32). Instead, over the time-scales discussed in this work, we have observed only a robust positive, causal impact of proliferation rate on stem cell competition that can be attributed to a key attribute of organization of those stem cells, namely division-independent differentiation.

## EXPERIMENTAL PROCEDURES

### Multicolor Twin-spot Lineage Analysis

Two-day old adult females of genotype *yw hs-flp/yw; ubi-GFP FRT40A FRT42B ubi-RFP/tub-lacZ FRT40A FRT42B* were given a single mild heat shock (32C for 30 min) to induce a low frequency of recombination in FSCs and ovaries were dissected 3 days later. We showed previously that in these flies fewer than one in a hundred ovarioles produced recombinant FSC genotypes in the absence of heat-shock (23). Hence, we can be sure that virtually all recombination events occur shortly after heat-shock. In all experiments flies were maintained at room temperature by frequent passage on normal rich food supplemented with fresh wet yeast. Ovaries were dissected and stained for Fasciclin 3 (Fas3) and β-galactosidase. Germaria were imaged in z-intervals of 3 μm and ovarioles in intervals of 4 μm on a Zeiss LSM 700 confocal microscope.

### Image acquisition and processing

Ovarioles were imaged with a 63x 1.4 N.A lens on a Zeiss LSM 700 confocal microscope (Carl Zeiss). Zeiss Zen 2012 was used to acquire microscope images of 512x512 pixels at 12 bit depth with line averaging 2 and pixel dwell 3.15 μs. Germaria and initial egg chambers were acquired with xzy scaling of 0.198 μm, 0.198 μm, 3 μm and larger egg chambers were acquired at 0.5 zoom, xyz scaling 0.397 μm, 0.397 μm, 4 μm. The range indicator was used to set the appropriate laser intensity for each fluorophore such that the signal was in the linear range and neither under- or oversaturated. Zen was used to linearly adjust channel levels when a color was dim rather than setting the laser intensity higher and photobleaching the sample. The presence of a fluorescent signal in each color was determined by examining each channel individually in the collected image (see Fig.S2). Images were digitally zoomed in Zen using digital interpolation and exported as tifs using “Contents of Image Window.” Images were rotated in Adobe Photoshop CS5 to position the anterior of the germarium and egg chambers to the left.

### Lineage analysis of mutant genotypes by MARCM

Flies with alleles on an *FRT40A* chromosome; *NM* (*Notch-Myc*, control), *cycE^WX^*, *cutlet^4.5.43^, pten^c494^*, with or without *UAS-cycE* on the 3^rd^ chromosome, were crossed to *hs-flp, UAS-GFP, tub-GAL4; tub-GAL80 FRT40A/Cyo; act>CD2>Gal4* flies for positive MARCM marking (“>” indicates an *FRT*). Flies with alleles on an *FRT42D* chromosome; control, *yki^B5^* and *hpo^42-47^* were crossed to *hs-flp, UAS-GFP, tub-GAL4; FRT42D tub-GAL80/Cyo; act>CD2>Gal4* flies for positive MARCM marking. Flies were dissected at 6d (after EdU labeling; see below) and 12d after heat-shock (30 min at 37C), and stained with antibodies to GFP and Fas3. FSCs, ECs, and FCs were scored in all z-sections of ovarioles.

### EdU Labeling

Ovaries were dissected directly into 15 μM EdU in Schneiders insect medium, incubated 1 h at room temperature and then fixed for 10 min at room temperature in 4% paraformaldehyde prior to antibody staining. EdU incorporation was detected using the Click-iT Plus EdU Imaging Kit C1063B (Life Technologies).

### Immunohistochemistry

Ovaries were fixed in 4% paraformaldehyde in PBS for 10 min at room temperature, rinsed three times in PBS with 0.1% Triton and 0.05% Tween-20 (PBST), and blocked in 10% normal goat serum (Jackson ImmunoResearch Laboratories, Inc.) in PBST. Monoclonal antibodies to Fasciclin III and Vasa were obtained from the Developmental Studies Hybridoma Bank, created by the NICHD of the NIH and maintained at The University of Iowa, Department of Biology, Iowa City, IA 52242. 7G10 anti-Fasciclin III was deposited to the DSHB by Goodman, C. and was used at 1:300 for multicolour lineage experiments and at 1:250 in all other experiments. Anti-Vasa (used at 1:10) was deposited by Spradling, A.C. / Williams, D. Other primary antibodies used were anti-β-galactosidase (Catalog No. 55976, MP Biomedicals) at 1:1000; anti-GFP (A6455, Molecular Probes). Ovarioles were incubated in primary antibody for 45 min for multicolour lineage experiments and overnight for all other experiments. Ovarioles were rinsed in PBST three times and incubated 1 h in secondary antibodies Alexa-488, Alexa-546, Alexa-594, or Alexa 647 from Molecular Probes. Alexa-546 and Alexa-647 were used in the multicolour lineage experiments to label Fasciclin III and β-galactosidase, respectively. Ovarioles from multicolor experiments were mounted in Prolong Gold Antifade (Invitrogen). DAPI-Fluoromount-G (Southern Biotech) was used as mounting medium for all other experiments.

## AUTHOR CONTRIBUTIONS

Conceptualization, A.R., D.M., and D.K.; Methodology, A.R., D.M., and D.K.; Formal Analysis, A.R., D.M. and D.K.; Investigation, A.R., and D.M.; Mathematical modeling, S.T.; Writing-Original Draft, D.K.; Writing-Review & Editing, A.R., D.M., and D.K.; Visualization, A.R. and D.K.; Funding Acquisition, D.K.

## ACKNOWLEDGMENTS

We thank Sarah Finkelstein for technical assistance, the Developmental Studies and the Bloomington Stock Center for antibodies and fly stocks, Tulle Hazelrigg, Alice Heicklen and Jamie Little for comments on the manuscript. This work was supported by the National Institutes of Health (RO1 GM079351 to D.K.); D.M. was supported in part by an NIH training grant.

